# Neuroendocrine control of catch-up growth in Drosophila

**DOI:** 10.1101/2022.12.30.522288

**Authors:** Diana M Vallejo, Ernesto Saez, Lucia García-López, Roberto Santoro, Maria Dominguez

## Abstract

Children and other vertebrate animals stunted due to malnutrition can compensate for this deficit by resuming growth at a higher-than-normal rate via a still ill-defined mechanism. High mortality and adverse effects later in life may offset the positive effects of catch-up growth. Here we report that the invertebrate *Drosophila melanogaster* also experiences catch-up growth following a period of starvation, and the relaxin receptor Lgr4 instigates this catch-up growth. Starved larvae compensate for weight loss by growing two or more times faster and starting maturation within the same time as the non-starved sibling by preventing a rise in insulin-like growth (IGF)-induced ecdysone under Lgr4 control. Our data reveal that catch-up growth is associated with a surge of insulin, not IGF, which may clarify how catch-up growth often leads to metabolic problems and obesity.

## Introduction

Growth is a universal process of life, and animal use thrift strategies to self-stabilize it in the face of threats such as poor nutrition, illness, or a local injury ^1,2^. Prader et al. (1963) ^3^ coined the term “catch-up” growth to describe the period of supranormal growth rates that often follows a transient growth inhibition and compensate and stabilize the growth trajectory. Nonetheless, not all children undergo catch-up growth, and those that catch up may experience adverse health problems later in life, including obesity and metabolic diseases ^3–5^. In addition to children ^6^, catch-up growth has been documented in most vertebrate groups, including other mammals ^7,8^, birds ^9^, reptiles ^10,11^, amphibians ^12^, and fishes ^13^. Despite its pervasiveness, the fundamental factor(s) and anatomical site involved in stimulating and limiting catch up growth are unknown ^14^, making effective interventions to reduce stunting and its side effects difficult ^5,15^.

Invertebrates such as fruit flies are excellent for investigating the genetic basis of developmental canalization ^16^ due to their experimental tractability and the documented ability of insect larvae to self-stabilize their size upon refeeding after a preceding period of growth retardation due to starvation ^17,18^. Here we use *D. melanogaster* third instar larvae to determine how juvenile organisms may recover their growth after a growth stunted due to starvation. We and others have recently shown that the brain, through the relaxin receptor Lgr3, is central for size stabilization in response to a local injury, tumour, or organ growth retardation caused by mutations ^19–24^. This discovery spurred our efforts to characterize the second relaxin receptor, Lgr4 ^25^ (Figure 1A), to investigate how refeeding allows recovery from stunting due to starvation. Specifically, we addressed the following questions: Is catch-up growth a universal phenomenon to compensate for a body growth deficit caused by starvation? Could the variability in recovery have a genetic basis? Finally, how does growth resume at a higher rate after starvation than the fed sibling’s growth rate?

**Figure 1.**
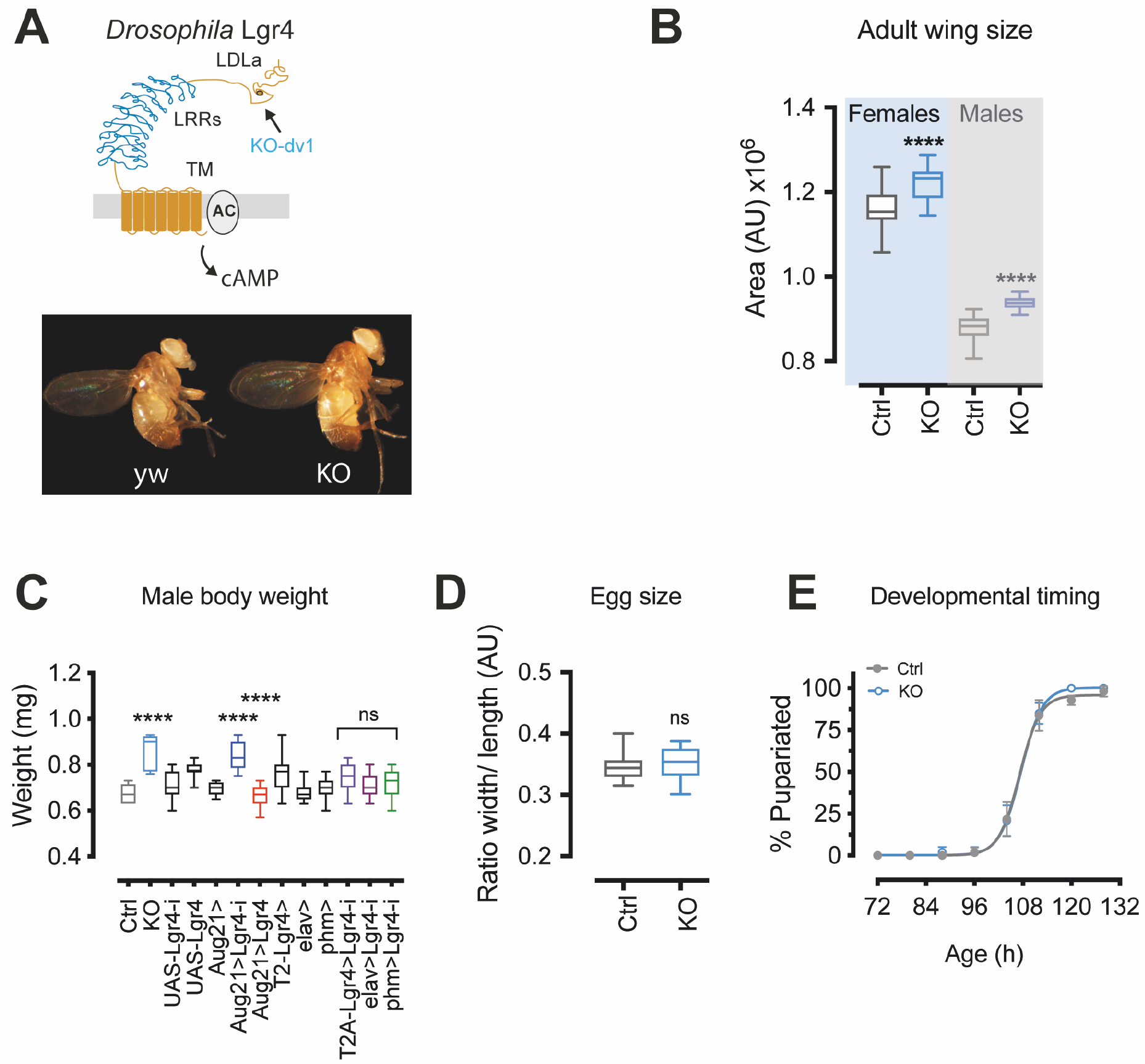
Relaxin Lgr4 acts in the corpus allatum and regulate systemic growth rate uncoupled of maturation time. (**A**) Lgr4 relaxin receptor and KO-dv1 mutation. Below are male flies of the *yw* CRISPR control and hemizygotes for the *Lgr4*^*KO-dv1*^ null allele (abbreviated KO). (**B**) Female and male adult wing size quantification of control and KO animals. (**C**) Adult male body size (weight). Control (Ctrl, *yw*) and *Lgr4*^*KO*^ (KO) and males with *Lgr4* silenced via RNAi (UAS-Lgr4-i) or overexpressed (UAS-Lgr4) tissue-specifically. *Aug21-Gal4* (*corpus allatum*), *phm-Gal4* (prothoracic gland), neurons (via *elav-Gal4*) and subsets of neurons (*T2-Lgr4-Gal4*). *n* ≥ 25 flies per genotype. (**D**) The eggs (*n*=20) derived from homozygous *Lgr4*^*KO*^ females and (**E**) developmental time to pupation formation (*n* = 60) are equal to those of control flies. Eggs size was measured as the width-to-length ratio. ^****^*P* < 0.0001, One-way ANOVA (**C** and **D**) and Unpaired *t*-test (**E**).

## Results

Towards these goals, we have generated a null allele of the relaxin receptor Lgr4 and an *UAS-Lgr4* transgene (Methods) for tissue-specific overexpression. Lgr4 relaxin receptor belongs to the LGR subfamily of GPCRs ^25,26^ and is homologous to the human Relaxin Family Peptide receptors RXFP1 (LGR7) and RXFP2 (LGR8) ^27^. Relaxin receptors have a unique leucine-rich repeat (LRR) essential for their activity ^28^. Therefore, we targeted this amino (N) terminal domain using the CRISPR/Cas9 technology producing null alleles of *Lgr4*. We backcrossed for ten generations of flies with null *Lgr4* mutations (hereafter *Lgr4*^*KO*^) and the control *yw* flies in which the CRISPR alleles were induced (hereafter *Ctrl*) before initiating the analyses.

Homozygous *Lgr4*^*KO*^ flies are viable and can be propagated the same way as wild-type sibling flies; however, some intergenerational vulnerability or trade-off is apparent and will be described elsewhere. *Lgr4*^*KO*^ flies are ostensible larger with proportionally larger structures (Figure 1, A and B). For example, the body size is 19% larger in males and 13 % in females compared to controls (Figure 1B and data not shown). We have used this trait to identify cells or anatomical loci where Lgr4 is required for systemic growth control. Surprisingly, knockdown of *Lgr4* with a validated RNAi transgene ^19^ in the *corpus allatum* (using the *Aug21-*Gal4 line ^29^) recapitulated the effect of wholly mutant *Lgr4*^*KO*^ (Figure 1C, and data not shown). Consistently, the overexpression of *Lgr4* in the *corpus allatum* produced the opposite effect, a reduction of final body size (Figure 1C). We verified by fluorescent in situ hybridization expression of *Lgr4* in the endocrine ring gland (data not shown). For simplicity, we refer to the specific overexpression in the *corpus allatum* gland as *Lgr4*^*OE*^. Ages of the larvae refer to hours after egg-laying.

Larger body size could reflect an increasing larval growth rate or growing time by postponing sexual maturation ^30^. Additionally, genetically larger mothers may lay larger eggs that develop into larger adults without changing growth rate or development time ^31^. We found that eggs laid by *Lgr4*^*KO*^ mothers were of equal size as those of control females (Figure 1D) and larva matured at the same age as control (Figure 1E) or even slightly earlier (data not shown). However, as shown below, compared with control larvae, *Lgr4*^*KO*^ larvae grow significantly faster (Figure 2B) while *Lgr4*^*OE*^ larvae grew slower (Figure 2C). Thus, the activity of the relaxin Lgr4 in the endocrine gland *corpus allatum* impacts the growth rate of well-fed larvae.

**Figure 2.**
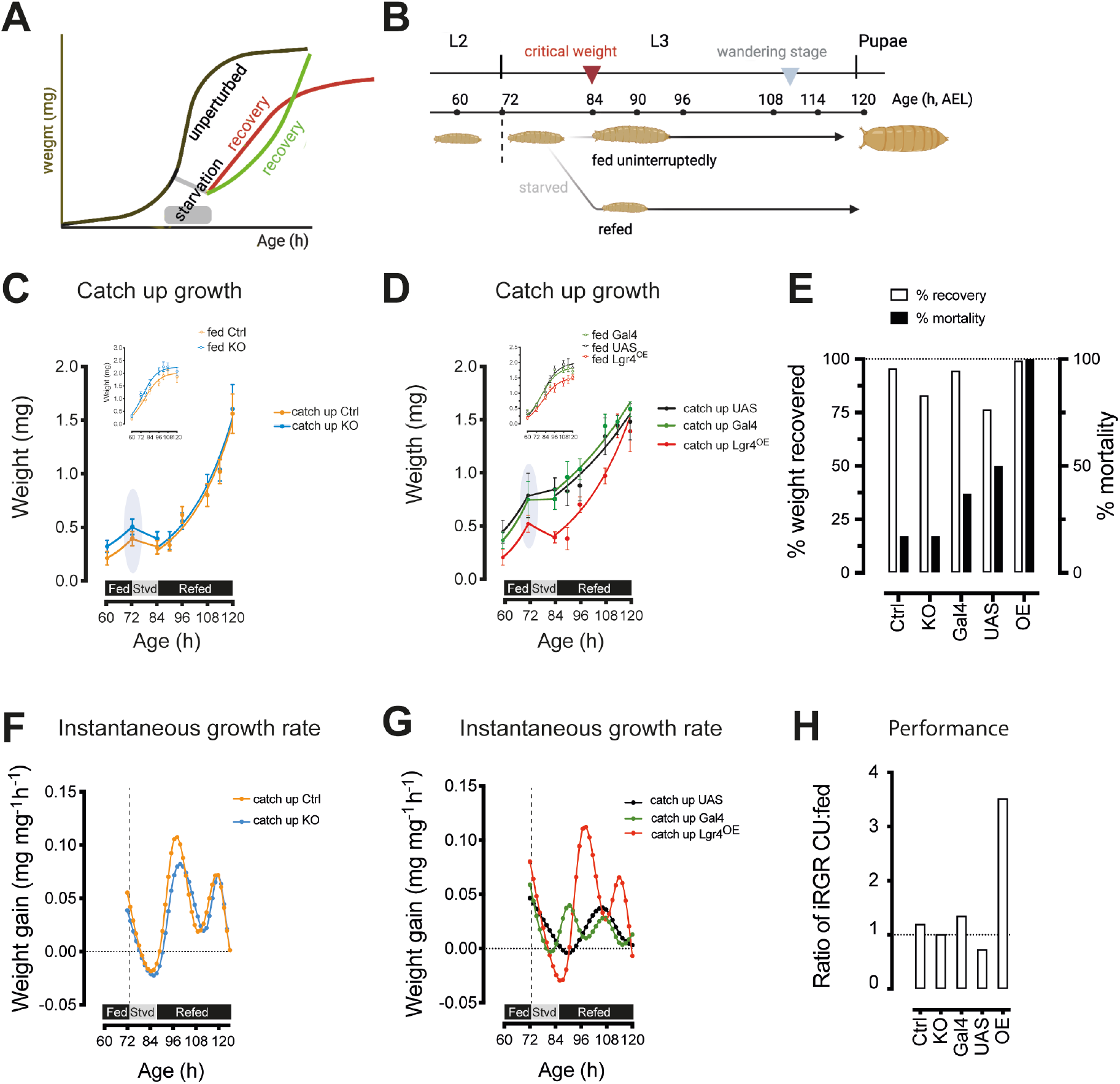
Catch up growth in Drosophila and stimulation by relaxin receptor Lgr4. (**A**) Refeeding after starvation may resume growth at a faster than normal rate (catch up growth, green line) or the normal rate (red line) but growing for a prolonged time to compensate for the previous growing deficit. (**B**) Top, timeline of insect developmental milestones during the transition from larva to pupa (hours after egg-laying, AEL). Bottom, starvation-refeeding regimen: Age-synchronized larvae were fed *ab libitum* until ecdysis to third instar larvae (L3) and then divided into two cohorts: one cohort was exposed to starvation for 12 hours at 72 h of age and then refed until voluntarily stopped feed and pupate, while the other was allowed to feed and grow without interruption. Body weight was measured at 6- or 12-hour intervals until pupation. (**C** and **D**) Growth curves of catch-up (CU) larvae and the unchallenged fed siblings (insets) from late L2 (60 h) until pupation. Each dot represents the average weight of 10 larvae, and error bars are the standard deviation (SD). Shading highlights body size at the initiation of starvation. Curve fitted non-linear regression (solid lines). The quality of the growth models was assessed using R^2^ and Akaike information criteria values (AICc) ^65^. Separate curves fit the three phases of growth during the fed-starvation-refeeding period. (**E**) Body weight recovery and mortality in each genotype. (**F** and **G**) The instantaneous relative growth rate for catch-up animals using restricted cubic regression splines with 95% confidence limits (Table S5). (**H**) The ratio of the relative growth rate of the CU period and weight gain of the fed sibling during the same period.

Animals may compensate for a growth deficit by accelerating the growth velocity (catch-up growth) ^6^ or growing for longer, postponing reproductive maturation (Figure 2A). For catch-up growth to occur, clinical and biological data show that the preceding period of starvation or illness that causes growth faltering must arrest maturation and that an excessively prolonged period of poor nutrition hinders recovery ^13,32^. To study whether catch-up growth occurs in insect larvae, we constructed growth curves of control larvae and Lgr4 mutant larvae during the starvation recovery. We examined instar-specific compensatory growth to avoid confusion due to the periods of stasis between moults. Specifically, we starved larvae shortly after they entered the last (third) instar stage (Figure 2B) and before they attained the critical weight ^33,34^, a developmental point after which larvae are irreversible commit the last instar larvae to metamorphosis and do not arrest their growth even after complete starvation^17,18^.

Larvae can withstand the complete absence of food for two days; however, after 24 h, they become very sluggish and sick ^17^. In contrast, 12-h starvation (with water) at 72 h of age effectively arrests growth and causes significant weight loss (up to 22% of pre-starvation weight) without causing immediate death. Importantly, during this interval, the larvae that continue to “fed” almost doubled their body mass (Figure 2, B and C) ^17^ due to the rapid growth of third instar larvae ^34^. Thus, larvae after starvation have a weight that is, on average, 50% or less than the expected weight of their chronologically aged and well-fed sibling.

Starved precritical weight L3 animals were allowed *ad libitum* feeding until they voluntarily stopped feeding and pupated (Figure 2B). Surprisingly, in all groups, starved-then-refed larvae entered the wandering stage and pupated at the same time as the continued-fed sibling (120 h, Figure 2, C and D). Thus, early third instar larvae can adjust growth rates to hasten recovery without postponing maturity, like observed in vertebrates. From now on, we refer to starved-then refed animals as “catch-up” animals.

The growth trajectories of animals that “catch-up” significantly differed from larvae of the same chronological age fed continuously (Figure 2, C and D). While third instar larval growth follows a typical S-shape (asymptotic) growth function ^35^, the growth of “catch-up” animals follow a J-shape (exponential) growth (curve fitting using exponential Malthusian repression function, Ctrl R^2^=89.67% and *Lgr4*^*KO*^ R^2^= 84.45%) as reported in children ^36^. The growth trajectory of “catch-up” *Lgr4*^*KO*^ larvae and its control did not differ (*F*_1.061_ = 7.7, *P* = 0.9397, unpaired t-test). However, their growth potential is very different. *Lgr4*^*KO*^fed larvae weigh 16% more than control larvae pre-starvation (shaded area in Figure 2C), so to a total return to the “fed” target size, *Lgr4*^*KO*^ larvae would need to grow faster than the control. Thus, *Lgr4*^*KO*^animals lose their growth advantage after being exposed to a period of starvation (Figure 2C). Incomplete recovery also occurred in the *w*^*1118*^ background in larvae with *corpus allatum*-specific silencing of *Lgr4* (w^1118^; *CA>Lgr4i*; 82.29% weight recovery). Conversely, *Lgr4*^*OE*^ achieved an impressive “catch-up” growth (99.30%, recovery Figure 2, D and E).

To put this catch-up growth into perspective, the maximum velocity peak in the “catch-up” Lgr4^OE^ larvae was 0.0783 ± 0.022 mg h^-1^ compared to the 0.047 ± 0.018 mg h^-1^ of the maximum velocity of the fed undisturbed sibling. The relative average instantaneous relative growth also provided an estimate of performance. For example, *Lgr4*^*OE*^ larvae recovered in only 24 h of refeeding (from 90 h to 114 h) on average 1.057 mg at an average speed of 4.72 ± 0.034 μg mg^-1^ h^-1^, while in the same period, the unchallenged *Lgr4*^*OE*^ larvae gained on average 0.38 mg at a mean velocity of 2.13 ± 0.021 μg mg^-1^ h^-1^. Together, these data disclose that insects use a “catch-up” growth strategy with features similar to those described in children ^14^. Stunted animals seem to match their velocity according to the growing deficit they experience, not the length or severity of malnutrition. Recovery was clearly genetically influenced (compare Figure 2D). Together, our findings support catch-up growth involving a mechanism that compares the current size with the expected size for age, as suggested more than five decades ago ^6^.

The accelerated growth after a preceding starvation was associated with animal mortality (Figure 2E), which varied in the different groups but was somewhat proportional to the effort of catching up, except in the *UAS-Lgr4* larvae. The death occurred during refeeding, beginning at 96-108 h (i.e., after 6-12 hours of refeeding). The high mortality of “catch-up” *Lgr4*^*OE*^ animals likely reflects the physiological cost of their accelerated growth or irreversible damage incurred by starvation. However, *Lgr4*^*OE*^ animals can endure two days of complete starvation like control without dying. Moreover, starvation without re-feeding in older larvae led to small yet viable adult *Lgr4*^*OE*^ flies that were also fertile. High mortality associated with catch-up is a widespread phenomenon seen in children and other species ^37^ and due, in part, to an ill-defined refeeding syndrome ^38^.

Sexual maturation is signalled by a pulse of the sexual maturation steroid hormone ecdysone ^34^. Starvation after critical weight increases ecdysone levels associated with accelerated maturation and earlier cessation of growth ^39^, which may hinder an animal’s ability for a full recovery. We thus wanted to determine whether accelerated growth rates or refeeding after starvation may affect ecdysone production. To this aim, we measured the expression of *phantom* (*phm*) (Figure 3), a critical step in ecdysone biosynthesis^31^.

**Figure 3.**
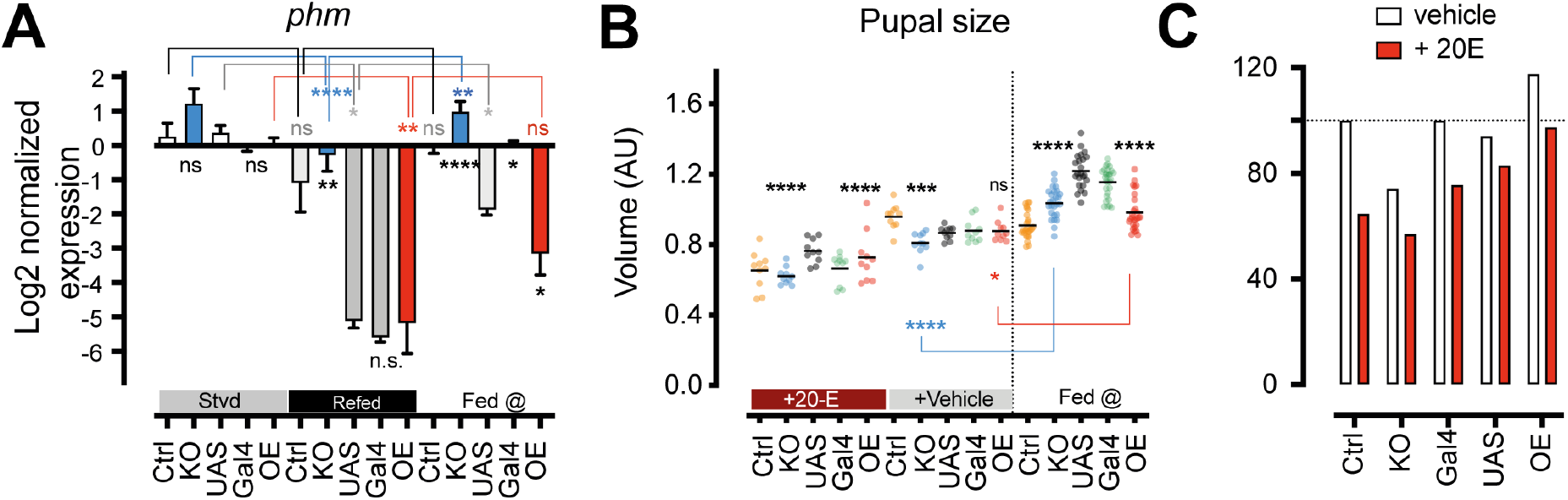
Ecdysone hinders catch up growth in *D. melanogaster*. (**A**) Levels of ecdysone biosynthetic enzyme *phm* measured by qRT-PCR. All larvae were at 96-102 h AEL. Strv, (24 h starvation starting at 72 h AEL); Refed (6 h refeeding after 24 h starvation); Fed (uninterrupted feeding). * P<0.1, ** P<0.01, *** P<0.001, **** P<0.0001, ns, no significant. Unpaired **t-test** and one-way ANOVA for multiple comparisons. The asterisks above denote statistical comparisons of the same genotype among nutritional regimens. The asterisks above are the comparisons within the same nutritional regimen. (**B**) Pupal size attained by control and mutant larvae refed with ecdysone (20E) or vehicle and compared to uninterrupted fed siblings. **(C)** Performance was measured as the ratio of weight gained of catch-up animal related to fed sibling.

While starvation for 24 h of 72 h old larvae did not change basal levels of ecdysone synthesis consistent with earlier studies (*30*) (Figure 3A), refeeding (6 h of feeding) caused a significant downregulation of ecdysone synthesis as compared with fed-unperturbed animals of the same chronological age (Figure 3A). *Lgr4*^*KO*^ larvae suffered a more pronounced downregulation of *phm* than control animals and animals with overexpressed Lgr4. Furthermore, we observed that *phm* levels changed, in the opposite direction, in larvae with loss and gain of *Lgr4*, meaning that relaxin receptor Lgr4 may somewhat counter the tendency of fast-growth larvae to produce the maturation hormone, thereby preventing precocious cessation of growth.

To test the role of ecdysone in recovery, we used the same groups and protocol as above but added 0.5 mg/ml of 20-hydroxyecdyone (20E, the active for ecdysone) during refeeding. We subsequently compared the performance (pupal size) by catch-up animals in relation to the performance of challenged animals without hormone treatment and the unchallenged animals (Figure 3B). In all groups, ecdysone treatment caused a profound impairment of recovery (Figure 3C), advancing that suppression of the steroid hormones may represent a mechanism for priming catch up growth.

These observations support a neuroendocrine control of catch-up growth, as proposed by Tanner in 1963 ^6^. Early studies often mistaken catch-up growth with the adolescence growth spurt, which also involves accelerating growth velocity ^40^. However, as observed by Tanners and other, the growth curves of catch-up children and adolescents are quite different ^6^. Similar, the catch up grow of catch-up insect larvae —J-shaped curves and those of mid-L3 larvae —S-shaped grow curve (Figure 2, C and D), suggest that catch-up grow and physiological growth spurts may be different.

To explore the mechanisms of catch-up growth, we next determined whether starvation and refeeding may modulate Lgr4 signalling to indicate how growth speed increases after refeeding following a period of growth retardation. To this aim, we measured the *corpus allatum* cyclic AMP (cAMP) (Methods), the main intracellular mediator of relaxin receptors ^19,22,41^. We used a cAMP Response Element (CRE>Luc) reporter construct ^19,42^ and measured luciferase levels using qRT-PCR. We found that nutritional deprivation of 72 h larvae (complete starvation, 24 h) dampened *corpora allata* cAMP (Figure 4A), resulting in levels of cAMP equivalent to those observed in fed larvae with *corpus allatum-* specific silencing of *Lgr4* (Figure 4A, Methods). Importantly, refeeding (6 h) resulted in rapid and magnified cAMP levels reactivation (Figure 4A). Thus, nutrition directly controls Lgr4 relaxin signalling activity to modulate growth rates. Moreover, Lgr4-cAMP ‘overshooting’ explains how starved-and-then-refed larvae might grow faster than the fed undisturbed control and why larvae with overexpressed *Lgr4* (e.g., *Lgr4*^*OE*^) experience a more intense catch-up growth than larvae with less or none *Lgr4*.

**Figure 4.**
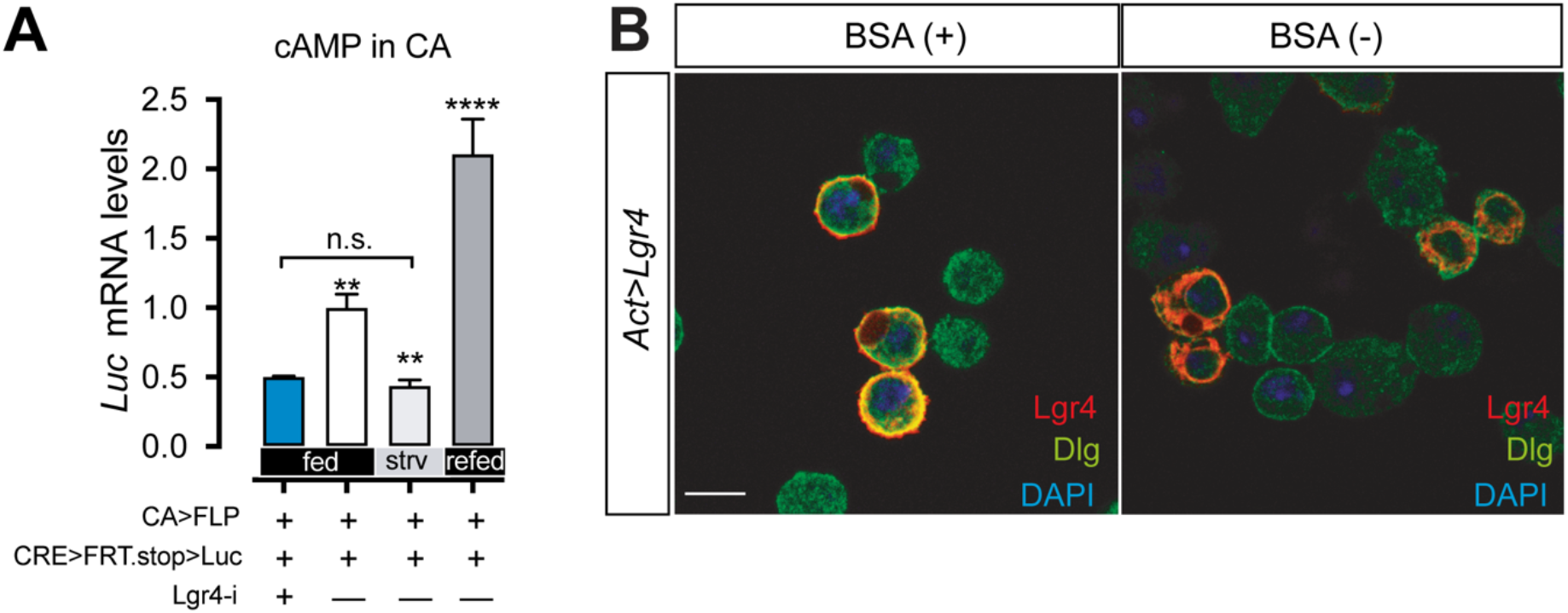
Nutritional control of Lgr4 activity. **(A)** Cyclic AMP responses of *corpus allatum* cells *in vivo* expressing the CRE>stop>Luc and the Flipase recombinase (UAS-Flp) or UAS-Lgr4-RNAi (Lgr4-i) under control of corpus allatum (CA) Aug21-Gal4 line. Experimental larvae were exposed to starvation for 24 hours or fed continuously. Responses were measured by transcription levels of luciferase and normalised against the housekeeping gene RP49. Data are shown as mean ± SD (n=10) and three replicated. (**B**) Serum starvation blocks traffic of Lgr4 protein to the plasma membrane. Scale bar, 10 μm. Confocal sections of Kc 157 cells were stained with anti-Lgr4 (red), anti-Dlg (green) and DAPI (blue).

Activating intracellular cAMP signalling by glucose and amino acids represents a vital nutrient-sensing system in many species ^27,43,44^. GPCRs can sense environmental fluctuations in nutrients directly. For example, receptor-trafficking, internalization and degradation regulate GPCR-signalling, and nutritional cues can directly regulate the activity of the LGR5 receptor in the murine intestine by regulating its subcellular localization ^45^. Traffic to the cell surface also regulates activation of murine relaxin receptors LGR7 and LGR8 (also called RXFP1 and RXFP2). We thus investigated the impact of starvation using an Lgr4 antibody (see Methods) to visualize the subcellular localization of the Lgr4 receptor in *Drosophila Kc167* cells transiently transfected with an *Lgr4* cDNA expression construct.

While in cells grown in the presence of foetal bovine serum, the Lgr4 receptor was present on the cell surface (Figure 4B), serum-starved cells transfected with the *Lgr4* cDNA showed cytoplasmic localization of Lgr4 and markedly reduction or complete absence of cell surface expression (Figure 4B). These data establish that Lgr4 relaxin signalling is nutritionally regulated, probably directly in the corpus allatum. Moreover, relaxin Lgr4 activity is magnified by refeeding following a period of starvation, consistent with Lgr4 effects on catch up growth.

A neuroendocrine hypothesis has long been proposed to explain the mechanism of catch-up growth ^14^; however, experimental data are still lacking ^4,40^. The Juvenile hormone (JH) is synthesized by the *corpus allatum*^46,47^ and plays a central role in larval growth, metabolism, reproduction, and longevity ^48,49^. To measure production and signalling by the JH, we monitored the expression of the Juvenile hormone acid methyltransferase (*Jhamt*) gene (data not shown), the rate-limited and final step in the JH biosynthesis pathway ^49–51^and *krüppel-homolog 1* (*kr-h1*; Figure 5A), the direct target and critical mediator of all juvenoids hormones and a well-documented surrogate of JH activity ^47,52^. Larval growth is also stimulated by insulin-like and IGF-like signalling ^31^. We measured the expression of FoxO target genes, *InR* and *4e-bp* ^53^, and levels of expression of *Ilp2* and *Ilp6* genes ^54^.

**Figure 5.**
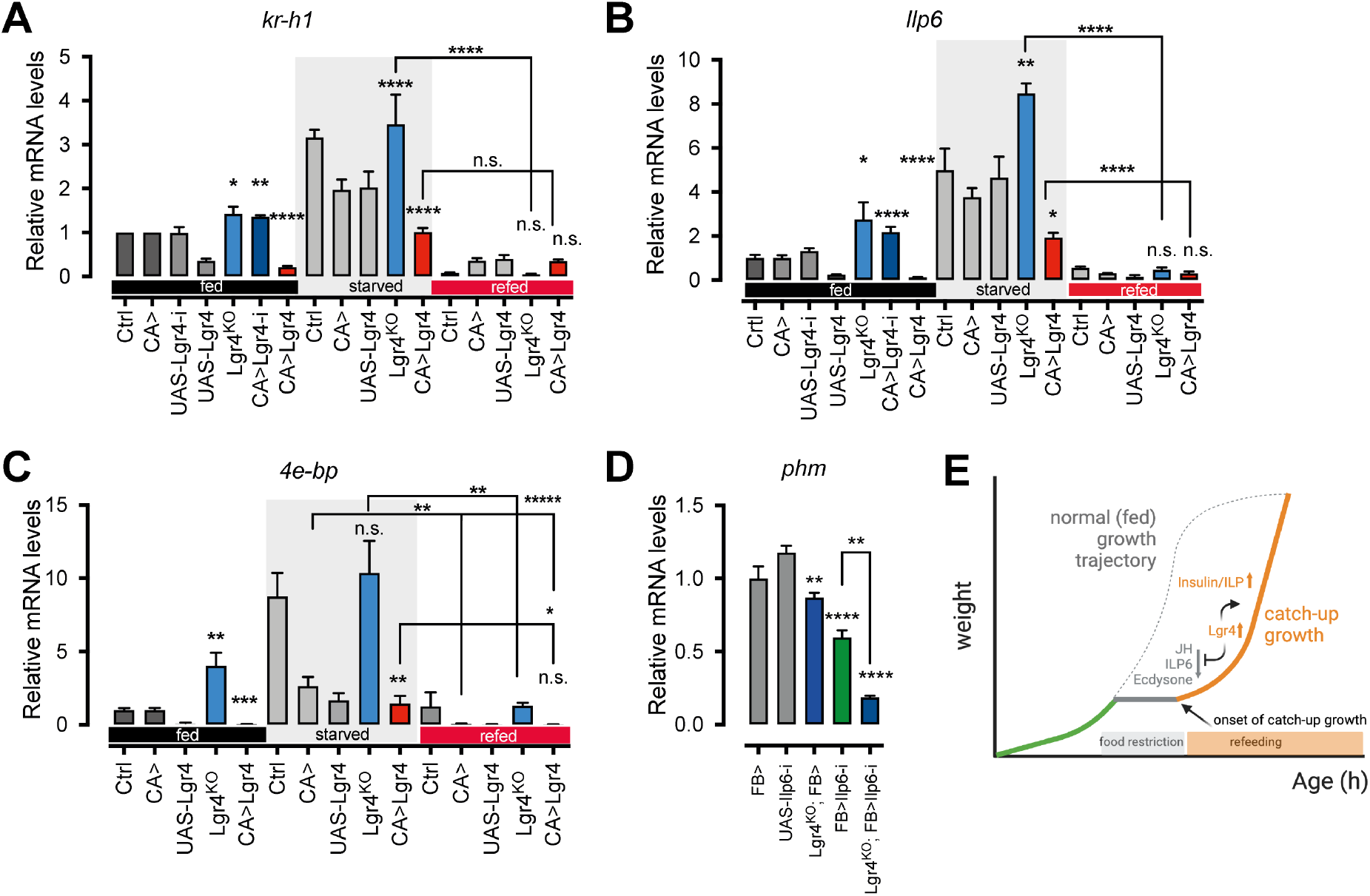
Starvation represses and refeeding restores insulin signalling and suppresses JH, ILP6 and ecdysone signalling. (**A-C**) Graphs show normalised expression of indicated genes to undisturbed fed (fed @) control of 96 hours of age. Starved means expression after 24 hours of complete starvation from 72 to 96h after egg laying (AEL). Refed means expression at 6 hours of feeding at 100 hours of age. Gal4, Aug21-Gal4/+; UAS, UAS-Lgr4, KO, Lgr4^KO^, Lgr4-i, Aug21-Gal4 UAS-Lgr4-RNAi. (**D**) The graph shows normalised expression of *phm* at 96 h in fed larvae of the indicated genotypes. FB (Fat body), Cg-Gal4. (E) Model of endocrine regulations during catch-up growth. The dashed line indicates the trajectory of unperturbed, fed larvae. Starved (24 h starvation starting at 72 h AEL); Refed (6 h refeeding after 24 h starvation); Fed (uninterrupted feeding). * P<0.1, ** P<0.01, *** P<0.001, **** P<0.0001, n.s., no significant. Unpaired t-test and one-way ANOVA for multiple comparisons.

Starving early third instar larvae (72 h of age) led to a marked increase in *Jhamt* and *kr-h1* (e.g., see Figure 5A). This starvation increase of JH is analogous to an elevation of growth hormone (GH) in other species ^55^. The IGF-like gene, ILP6, was also upregulated during starvation (Figure 5B), and this starvation programme was abrogated entirely in starved *Lgr4*^*OE*^ larvae (Figure 5, A and B).

Studies of catch-up growth in fish suggested that elevated GH persistence could lead to an overstimulation of IGF-1 when feeding is restored to stimulate accelerated growth ^56^.4. This “carryover” model is attractive but lacks experimental evidence of overlapping expression of IGF-1 and catch-up growth, as seen during the physiological pubertal growth spurt ^14^. We found that refeeding (6 h) downregulated *kr-h1* and *Ilp6* rapidly and below levels of fed control (Figure 5, A and B), as seen with ecdysone (Figure 3). The transcriptional repressor 4E-BP/Thor is a surrogate of larval growth rate and a transcriptional target of the insulin pathway ^53^. Levels of 4E-BP/Thor were significant upregulation during starvation and rapidly downregulated after refeeding (Figure 5C), consistent with insulin/InR/FoxO signalling mediating catch up growth.

Downregulation of ecdysone upon refeeding when insulin increases suggests that biosynthesis of ecdysone may be under control IGF-1/ILP6, rather than insulin/ILP2. Therefore, we tested whether silencing endogenous *Ilp6* in the fat body ^57^ via the Cg-Gal4 may affect basal ecdysone. Indeed, the knockdown of *Ilp6* led to a drastic downregulation of *phm* (Figure 5D). Collectively, these findings show that relaxin Lgr4 indirectly affects ecdysone vi its regulation of ILP6, which facilitates catch-up growth via anabolic insulin signalling (Figure 5E).

## Discussion

Catch-up growth illustrates the resilience of juvenile organisms to body growth disruptions that potentially reduce their fitness. Intriguingly, self-stabilization of whole-body growth retardation and a local injury or organ growth inhibition entangles members of the same family, the relaxin family. However, the compensatory mechanisms are different, with distinct outcomes. For example, the presence of a local injury, or a mutation that inhibits the growth of an organ or body part, activates a developmental checkpoint via Lgr3 receptor signalling that leads to delayed maturation to ensure sufficient time to repair. This strategy corrects size but delays reproduction, increasing the risk of dying without offspring. Whole-body growth retardation caused by malnutrition, infections, or malabsorption-causing diseases, is typically compensated for by accelerating growth rate through a mechanism that decouples sexual maturation to avoid an advanced maturation time that would impede a complete body size recovery.

It is not difficult to imagine how malnutrition affects the organism’s growth, and food restoration allows it to resume. However, it is more problematic to envision how an exhausted animal, with depleted energy reserves and shrunken alimentary organs, can, when fed again, resume growth at a rate several times the growth rate of those fed continuously. We speculate that the struggle to achieve a higher-than-normal growth rate, not just the impact of starvation, is responsible for the high mortality and adverse long-term consequences of spontaneous catch-up growth. Moreover, an immediate goal of catch-up organisms may be to replenish the depleted internal resources as fast as possible, which may explain why gain in weight in children and other animals begins before height gain after food rehabilitation ^2,58^.

Stunting remains a significant health problem that hampers a child’s physical and cognitive performance and future health ^59^. Stunting is strongly interconnected to poverty and food insecurity ^60^, and often creates a cycle that stunted mothers and offspring cannot escape. Understanding the endocrine determinants of catch-up growth seems promising for leveraging those factors to promote fuller recovery and less adverse consequences. Our study advances that the onset of growth catch-up is related to the anabolic insulin to replenish energy stores rapidly and that children with a genetic predisposition for energy storage may have a greater ability to catch up yet a higher risk of obesity and other adverse health effects of spontaneous catch-up.

## Materials and Methods

### Drosophila stocks and husbandry

*Aug21-Gal4/CyO, Lgr4-RNAi*^*BL28655*^*(P*{*TRiP*.*JF03070*}*attP2), w*^*1118*^, *elav-Gal4 (P*{*GawB*}*elav[C155]), phm-Gal4, p206-Gal4, y*^*1*^ *w*^*\**^ *Mi*{*Trojan-GAL4*.*1*}*Lgr4*^*MI06794-TG4*^, *UAS-FLP, Cg-Gal4 (P*{*Cg-GAL4*.*A*}*2), dilp6-RNAi*^*BL33684*^*(P*{*TRiP*.*HMS00549*}*attP2)*, and *yw*, were obtained from the Bloomington Stock Centre at Indiana University. *CRE-F-Luc* was a gift from J. C. P. Yin ^42^.

*UAS-Lgr4* transgenic flies were generated by injection of the *pUASt-Lgr4* construct ^19^ in *w*^*1118*^ embryos following standard P-element-mediated transformation procedures (BestGene, InC).

All experimental crosses were performed at 26.5ºC in Nutri-Fly® food (Cat #: 66-112 Bloomington Formulation, Genesee Scientific) on a 12:12-hour light: dark cycle.

### CRISPR/Cas9 Lgr4 allele generation

The *Lgr4* allele used in this study was generated using the CRISPR/Cas9 system ^61^. Two gRNAs targeting the exon 2 of *Lgr4* were designed and used to generate the following oligos to create a pCFD4 construct (pCFD4-U6:1_U6:3tandemgRNAs was a gift from Simon Bullock (Addgene plasmid # 49411):

Lgr4-CRISPR-forward:

5’TATATAGGAAAGATATCCGGGTGAACTTCG**<u>CGAGAGTAAATGTGATAGGC</u>**GTTTTAG AGCTAGAAATAGCAAG-3’

Lgr4-CRISPR-reverse:

5’ATTTTAACTTGCTATTTCTAGCTCTAAAAC**<u>TGCGCGCCTTCATTAGAAGC</u>**CGACGTTA AATTGAAAATAGGTC-3’

gRNA sequences targeting *Lgr4* are underlined and in bold. To introduce the protospacer sequences into pCDF4 we run a PCR using the Lgr4-CRISPR-forward and reverse primers and pCDF4 plasmid as template using the Thermo Scientific Phusion Hot Start II high-fidelity DNA polymerase (Cat #F549S, ThermoFisher Scientific). In parallel, pCDF4 was digested using BbsI restriction enzyme (Cat #R0539S New England Biolabs). PCR reaction and digested plasmid were gel purified and used to perform a Gibson Assembly reaction (Cat. #E2611S, New England Biolabs) followed by bacterial transformation according to the manufacturer instructions. Insert was verified by sequencing. Injections were performed by BestGene, in *yw nos-Cas9* embryos (BDSC #54591). To remove the *nos-Cas9* genetic background, the F0 adults that emerged were crossed to *y*^*1*^*w*^*1*^ flies. Males obtained were then crossed to *FM7i* balancer stock females. Progeny lacking the *nos-Cas9* transgene (lacking the *mini-white* marker) was then selected for establishing each of the CRISPR mutant candidate lines.

To identify CRISPR induced mutations, genomic DNA was obtained from each putative mutant line and used as template for PCR amplification of the Lgr4 target sequence using the following primers: 5’-GCCATAAGTTAGCCGAATGG-3’ and 5’-CTCGACAGGGACAGGAAAAG-3’.

PCR products were then verified by sequencing using the primer:

5’-GGCTTGTTAAATCGGCACTT-3’.

In this screen, we identified two alleles inducing a change in the open reading frame ORF of the Lgr4 protein that induced the formation of two premature stop codons in tandem in the exon 2. Both alleles are null and phenotypically identical and in this study the allele *Lgr4*^*KO-dv1*^ was chosen to perform all experiments.

### Adult weight and wing size measurements

For weighing adult flies, 20 females and 20 males were crossed and left 24 hours for egg deposition in Nutri-Fly® food. Parental flies were transferred every 24 hours to fresh tubes and laid eggs were reared at 26,5ºC. Eclosed males and females of each genotype were collected and incubated 24 hours at 26.5ºC. Flies were weighted individually three times using a precision scale. Final weight of each fly is the average of the three measurements.

For adult wing measurements, 20 females and 20 males were crossed and left 24 hours for egg deposition Nutri-Fly® food. Parental flies were transferred every 24 hours to fresh tubes and laid eggs were reared at 26.5ºC. Adult males and females were collected, and wings were excised and rinsed thoroughly with ethanol and mounted in a 70% glycerol solution in PBS. Wing areas were measured using ImageJ.

### Pupal volume measurement

For pupal volume determination, 20 females and 20 males were crossed and left 24 hours for egg deposition Nutri-Fly® food. Parental flies were transferred every 24 hours to fresh tubes and laid eggs were reared at 26,5ºC. Pupae were collected and photographed with their dorsal side up. Length and width were measured using ImageJ; volume was calculated according to the following formula: v = 4/3p(L/2)(l/2)2 (L, length; l, width).

### Time-course growth rate measurement

Females and males (20 to 30 of each) were crossed and, after 24 hours mating at 26.5°C, parents were transferred to grape juice agar plates with yeast paste and left 4 hours for egg deposition. Adults were removed, and laid eggs were incubated 24 hours at 26.5°C. First-instar eclosed larvae were transferred onto 5 ml-plates of fresh Nutri-Fly® food (30 larvae per plate). Larvae were harvested and carefully cleansed for any remaining food, and then weighted individually three times every 6 or 12 hours using a precision scale. Final weight of each larva is the average of the three measurements.

For time-course growth rate measurement in starvation and refeeding assays laid eggs were incubated 24 hours and first instar eclosed larvae were transferred onto 5 ml-plates of fresh Nutri-Fly® food (30 larvae per plate, in triplicates as minimum) and reared until ecdysis to L3 (∼for 36 hours at 26.5°C). L3 larvae were harvested and weighted as described above and at 72 hours after egg laying (AEL), they were transferred onto 2% agarose plates with water and incubated until 84 AEL. Starved larvae were then weighted and carefully transferred back onto fresh 5 ml-plates of Nutri-Fly® food. Larvae were then harvested and weighted every 6 or 12 hours. As post-starved larvae were significantly weak, we performed experiments in large cohorts and weighted different individuals for each time point (10 larvae in total per time point) to reduce any further damage of the animals during manipulation.

### Egg size measurement

Females and males that were up to three days old were collected and let mating in Nutri-Fly® food supplemented with yeast. After three days mating, females’ ovaries were dissected in cold PBS and fixed in 4% paraformaldehyde. Ovarioles were split up and stage-14 oocytes (mature eggs), identified for the elongation of the dorsal filaments, were mounted in Vectashield mounting medium (Cat. # H-1000, Vector Labs) maintaining their three-dimensional (3D) configuration ^62^ and photographed along the dorsoventral axis with their lateral side up. Individual egg length and width were measured using ImageJ.

### Ecdysone treatment

Age-synchronized larvae were collected and starved as described above. Starved larvae were transferred onto fresh Nutri-Fly® food (30 larvae per vial) supplemented with 20-hydroxyecdysone (20E) (Sigma H5142-10MG) in a concentration of 0.5 mg/ml. Vehicle control (ethanol) was performed in parallel. Larvae were allowed to feed *ad libitum* until they voluntarily exited the food and pupated. The pupal size was measured as described above.

### Characterization of Lgr4^KO^ allele in Drosophila cultured cells

*Drosophila Kc167* cells were obtained from the Drosophila Genomics Resource Center (DGRC). Cells were cultured in Schneider’s Drosophila medium (Cat. # 21720024, Invitrogen) supplemented with 10% Foetal Bovine Serum (Cat. # 10500064, Gibco) at 25°C in a non-humidified, ambient-air-regulated incubator.

*Kc167* cells (8×105 cells per well) were co-transfected in 6-well plates with *pActin-Gal4* and either *pUAS-Lgr4, pUAS-Lgr4KO-dv1* or empty *pUASt* vector using Fugene-HD (Cat. # E2311, Promega). Forty-eight hours after transfection, cells were washed twice with PBS, fixed using 4% paraformaldehyde and immunostained (cells were permeabilized using PBS + Triton 0.1%). Primary antibodies used: guinea pig anti-Lgr4 (1/100) and mouse anti-Dlg (1/100, 4F3 Developmental Studies Hybridoma Bank (DSHB)). Secondary antibodies were purchased from Invitrogen. Cells were mounted in Vectashield antifade mounting medium with DAPI (Cat. # H-1200, Vector Labs)

### Drosophila cultured cells starvation assay

*Kc167* cells (8×105 cells per well) were co-transfected in the presence of 10% Foetal Bovine Serum (Cat. # 10500064, Gibco) in 6-well plates with *pActin-Gal4* and *pUAS-Lgr4* vector using Fugene-HD (Cat. # E2311, Promega). Twenty-four hours after transfection, the serum-supplemented medium was removed and replaced by serum-free Schneider’s *Drosophila* medium. Cells were incubated 24 hours and then washed twice with PBS, fixed using 4% paraformaldehyde and immunostained (cells were permeabilized using PBS + Triton 0.1%). Primary antibodies used: guinea pig anti-Lgr4 (1/100) and mouse anti-Dlg (1/100, 4F3 Developmental Studies Hybridoma Bank (DSHB)). Secondary antibodies were purchased from Invitrogen. Cells were mounted in Vectashield mounting medium (Cat. # H-1000, Vector Labs).

### Lgr4 antibody

The extracellular domain fragment of Lgr4 (from 51 to 400 amino acids) was synthesized and used for the immunization of guinea pigs. Antigen synthesis, immunization and antibody purification were performed by Genscript Biotech Corporation (Piscataway, NJ, USA) according to their polyclonal antibody programme.

### Western Blot analysis

*Kc167* cells (8×105 cells per well) were co-transfected in 6-well plates with *pActin-Gal4* and *pUAS-Lgr4, pUAS-Lgr4KO-dv1*or *pUAS-Lgr4KO-dv2* vector using Fugene-HD (Cat. # E2311, Promega). Forty-eight hours after transfection, cells were washed twice with PBS and then lysed using modified RIPA buffer containing proteinase inhibitors. Protein samples were separated in a 4-12% SDS-PAGE gel (Cat. # XP04120BOX, Invitrogen) and transferred to a PVDF membrane (Cat. # IPVH00010, Millipore). The membrane was blocked and then incubated with polyclonal guinea pig anti-Lg4 (1/500). HRP-conjugated rabbit anti-Guinea Pig (Cat. # 61-4620, Thermo) was used as the secondary antibody. Protein was detected using the chemiluminescent substrate ECL (Cat. # 32209, Pierce) in an Amersham Imager 680 (GE Healthcare) detector.

### Larval starvation for mRNA expression analysis

Females and males (20 to 30 of each) were crossed and, after 24 hours mating, parentals were transferred to grape juice agar plates with yeast paste and left 4 hours for egg deposition. Adults were removed, and laid eggs were incubated 24 hours at 26.5°C. First-instar eclosed larvae were transferred onto fresh Nutri-Fly® food (30 larvae per vial) and reared at 26.5°C for 48h hours. Seventy-two hours AEL larvae were taken out of the food and transferred into 2% agar vials (10-15 larvae per vial). Larvae were incubated for 24 hours at 26.5°C and then collected and snap-frozen for further mRNA expression analysis.

### Triacylglycerol (TAG) and protein measurements

TAG, glucose and protein levels were measured as described previously, with minor modifications ^63^. Synchronized feeding larvae (96 hours AEL) were collected and rinsed several times with cold PBS to remove all traces of food and snap-frozen. Samples (2 larvae per tube) were homogenized in a TissueLyser-LT (Qiagen) in 200 μl of the corresponding buffer according to the assay, centrifuged and the supernatant transferred to a clean tube. For each sample, 10 μl were saved at −80ºC for posterior determination of protein content using Pierce BCA Protein Assay Kit (Cat. # 23227, Thermo Scientific). Samples were then immediately heat-inactivated and cooled down to room temperature.

TAG determination: 60 μl of each sample were mixed with 60 μl of Triglyceride Reagent (Cat. # T2449, Sigma-Aldrich), incubated at 37 °C for 60 min, and cleared by centrifugation. 30 μl of supernatant were mixed with 100 μl Free Glycerol Reagent (Cat. # F6428, Sigma-Aldrich) in a 96-well plate and incubated for 5 min at 37°C. The absorbance of each sample was measured at 540 nm in an EZ Read 400 Microplate Reader (Biochrom). Triolein Equivalent Glycerol Standard 2.5mg/ml (Cat. # G7793, Sigma Aldrich) was used as standard. TAG concentration was calculated according to the manufacturer’s instructions. Data were normalized against protein content.

### Quantitative RT-PCR

Total RNA was extracted from *Drosophila* larvae using the RNAeasy-Mini Kit (Cat. # 74106, Qiagen). cDNA was synthesized using SuperScript® III First-Strand Synthesis System (Cat. # 18418020, Invitrogen) using oligo-dT primers. Quantitative real-time PCR was performed using Power SYBR Green PCR Master Mix (Cat. # 4367659, Applied Biosystems) using gene-specific primers on an ABI7500 Real-time PCR (Applied Biosystems) or a Quant Studio 3 Real-Time PCR system (Applied Biosystems) apparatus. *Rp49* primers were used for mRNA normalization. Relative gene expression was assessed by the qPCR quantification of three biological replicates for each condition calculated using the comparative Ct method.

Primer list:

*Jhamt:*

Forward: 5′-ATTCGCATCGACCATGCAGT-3′

Reverse: 5′-GAAGTCCATGAGCACGTTACC-3′

*Kr-h1:*

Forward: 5′-ACAATTTTATGATTCAGCCACAACC-3′

Reverse: 5′-GTTAGTGGAGGCGGAACCTG-3′

*Ilp6:*

Forward: 5′-CCCTTGGCGATGTATTTCC-3′

Reverse: 5′-CACAAATCGGTTACGTTCTGC-3′

*Ilp2:*

Forward: 5′-GGTGCCGACAGCGATCTG -3′

Reverse: 5′-TTCGCAGCGGTTCCGATA-3′

*InR:*

Forward: 5′-GCTGTCAAGCAAGCAGTGAA-3′

Reverse: 5′-TCTTTTTACCCGTCGTCTCC-3′

*4e-bp:*

Forward: 5′-GAAGGTTGTCATCTCGGATCC-3′

Reverse: 5′-ATGAAAGCCCGCTCGTAG-3′

*phm:*

Forward: 5′-TAAAGGCCTTGGGCATGA-3′

Reverse: 5′-TTTGCCTCAGTATCGAAAAGC-3′

*Luciferase:*

Forward: 5′-AGGTCTTCCCGACGATGA-3′

Reverse: 5′-GTCTTTCCGTGCTCCAAAAC-3′

*Rp49:*

Forward 5′-TGTCCTTCCAGCTTCAAGATGACCATC-3′

Reverse 5′-CTTGGGCTTGCGCCATTTGTG-3′

### Measurement of the developmental timing of pupariation

Females and males (20 of each) were crossed and, after 24 hours mating, parents were transferred to grape juice agar plates with yeast paste and left 4 hours for egg deposition. Adults were removed, and laid eggs were incubated for 48 hours at 26.5°C. Second instar larvae were transferred onto fresh Nutri-Fly® food (20 larvae per vial) and reared at 26.5°C. A survey of the pupae was performed at 4 hours intervals using flyGear, a custom-made automated high-performance pupal counting device.

### Single-Molecule Fluorescence In Situ Hybridization (smFISH)

Single-molecule FISH was performed using Stellaris (Biosearch Technologies) oligonucleotide probes 20 nucleotides in length complementary to the intron transcript of the gene *Lgr4* (48 probes, listed in Sup. Table 1) conjugated to Quasar-570. Ring glands of 96h AEL *w*^*1118*^ feeding larvae were dissected and processed as previously described ^64^. Processed tissue was then mounted in Vectashield mounting medium (Cat. # H-1000, Vector Labs), maintaining its 3D configuration. Images were obtained with a Super-resolution Inverted Confocal Microscope Zeiss LSM 880-Airyscan Elyra PS.1.

The mFISH probes for *Lgr4* are:

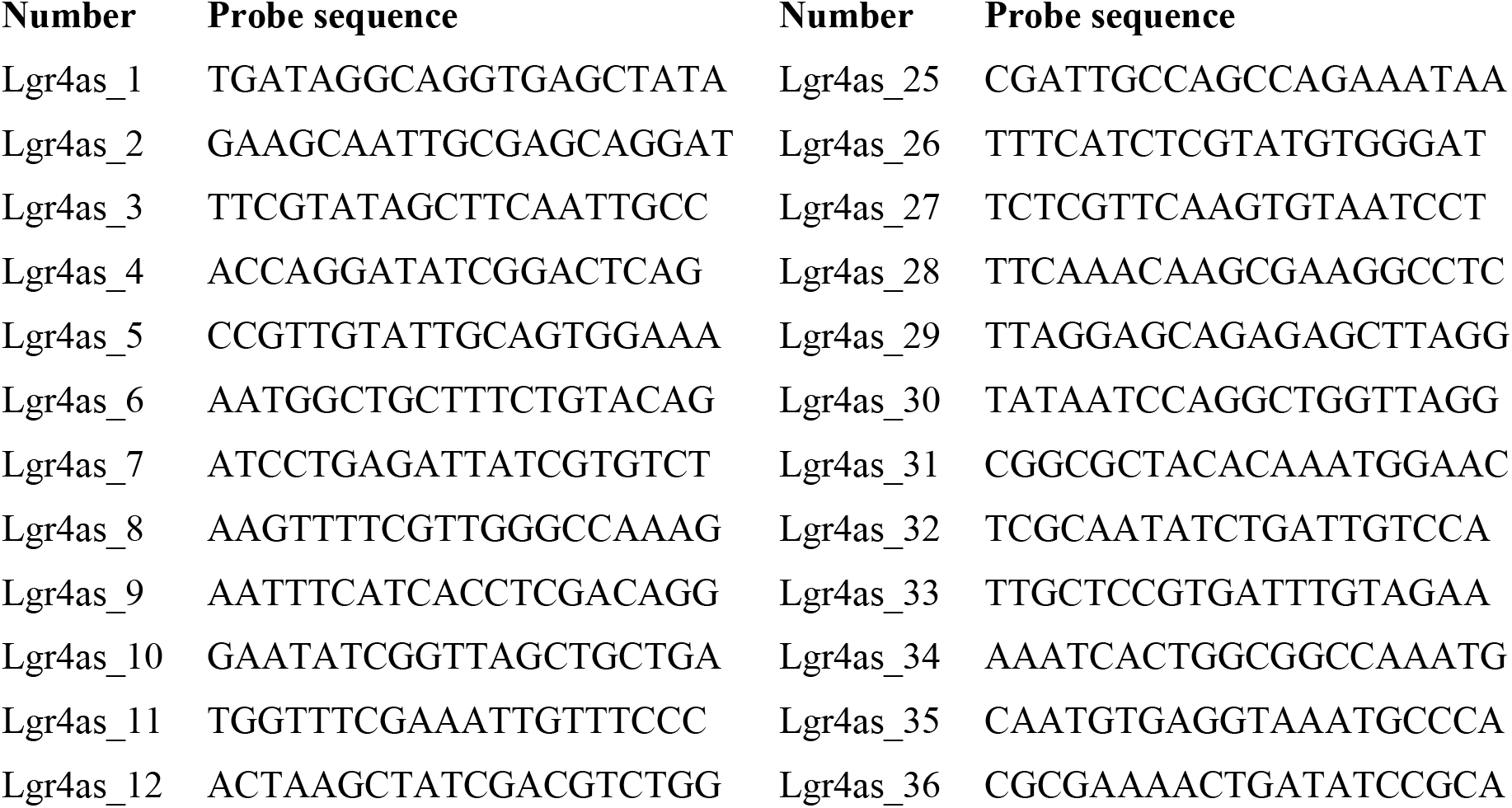
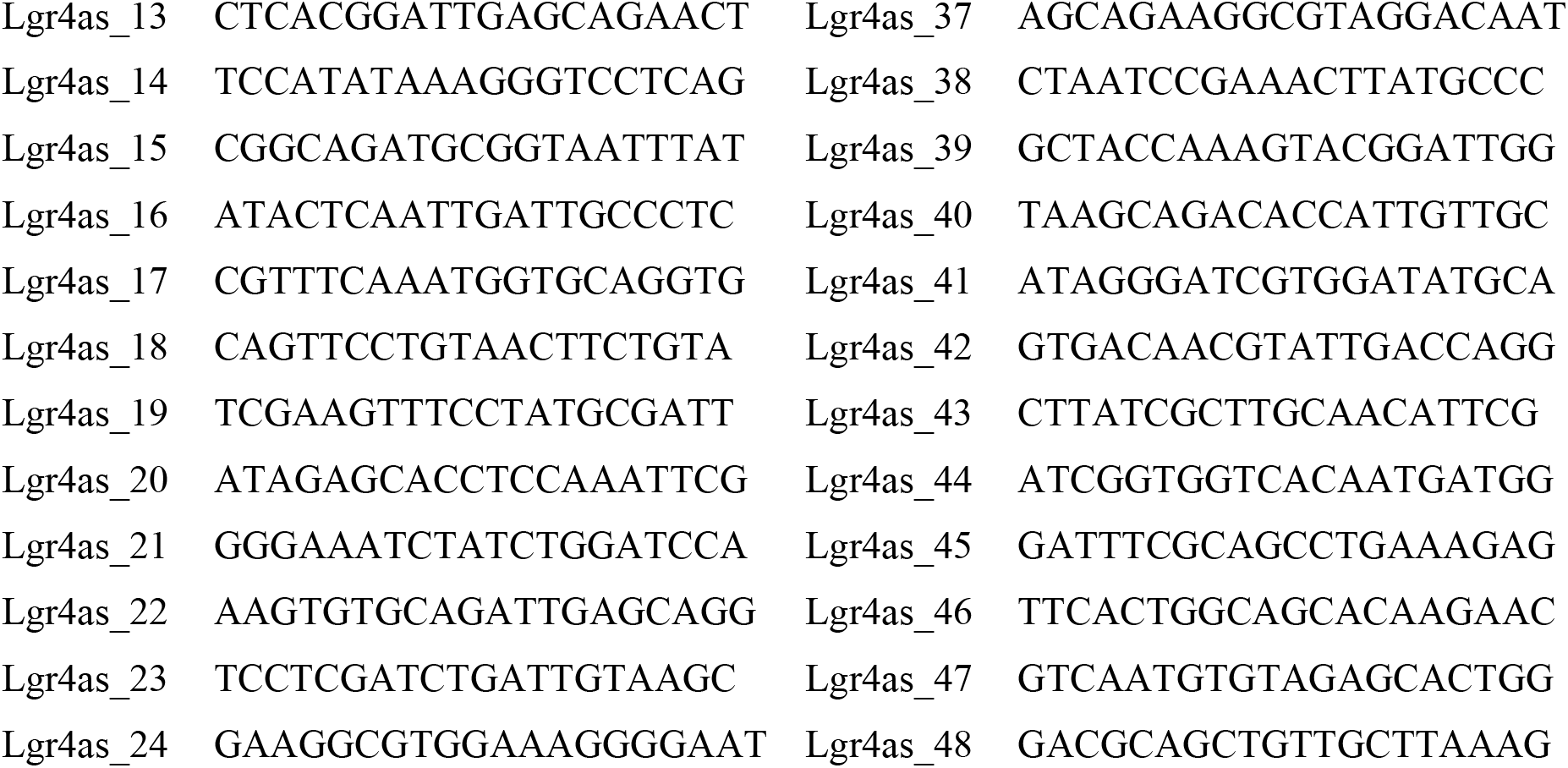

## Acknowledgments

We thank Laura Mira, Esther Ballesta-Illan, and Alicia Estirado for technical assistance. We also thank the Bloomington Stock Center (NIH <u>P40OD018537</u>), the TRiP at Harvard Medical School (NIH/NIGMS <u>R01-GM084947</u>), the Vienna Drosophila Stock Centre, for providing fly stocks, and the Developmental Studies Hybridoma Bank at the University of Iowa for antibodies. Figure 2A and 5E. were created with BioRender.com. We also thank G. Exposito and V. Villar for assistance with Airyscan microscopy. This work was supported by the Spanish National Grants (BFU2015-64239-R and PID2019-106002RB-I00 and PDC2022-13387-I00) funded by MCIN/AEI/10.13039/50000033 and co-financed by “ERDF A way of making Europe”, by the Fundación Científica Española Contra el Cáncer (AECC) (<u>CICPF16001DOMÍ</u>), and Generalitat Valenciana Grants (PROMETEO/2017/146 and PROMETEO/2021/027) to M.D. M.D. was also funded by the “Severo Ochoa” Program for Centres of Excellence in R&D (SEV-2017-0723 and CEX2021-001165-S) funded by MCIN/AEI/10.13039/50000033 and co-financed by “ERDF A way of making Europe”. E. S-C and R.S. are Spanish doctoral FPI fellows (PRE2018-085912 and BES-2016-077689) from the Spanish Ministerio de Ciencia e Innovación.

